# Few amino acid mutations in H6 influenza A virus from South American lineage increase pathogenicity in poultry

**DOI:** 10.1101/2022.04.28.489980

**Authors:** Agustina Rimondi, Valeria S. Olivera, Ivana Soria, Gustavo D. Parisi, Martin Rumbo, Daniel R. Perez

## Abstract

In chickens, infections with influenza A virus (IAV) can range from mild to severe and lethal. The study of IAV infections in poultry has been mostly limited to strains from the North American and Eurasian lineages, whereas limited information exists on similar studies with strains from the South American lineage (SAm). To better evaluate the risk of introduction of a prototypical SAm IAV strain into poultry, chickens were infected with a wild-type SAm origin strain (WT557/H6N2). The resulting virus progeny was serially passage in chickens 20 times and the immunopathological effects in chickens of the last passage virus, 20Ch557/H6N2, was compared to those of the parental strain. Comparison of complete viral genome sequences indicated that the 20Ch557/H6N2 strain contained 13 amino acid differences compared to the wild type strain. Five of these mutations are in functionally relevant regions of the viral surface glycoproteins hemagglutinin (HA) and neuraminidase (NA). However, despite higher and more prolonged virus shedding in chickens inoculated with the 20Ch557/H6N2 strain compared to those that received the WT557/H6N2 strain, transmission to naïve chickens was not observed for either group. Analyses by flow cytometry of mononuclear cells and lymphocyte subpopulations from the lamina propria and intraepithelial lymphocytic cells (IELs) from ileum revealed significant increase in the percentages of CD3+TCRγδ+ IELs in chickens inoculated with the 20Ch557/H6N2 strain compared to those inoculated with the WT557/H6N2 strain.

## INTRODUCTION

IAVs are enveloped viruses in the family Orthomyxoviridae containing a segmented negative sense single stranded RNA that encodes for 10 major ORFs and additional 2 to 4 minor ORFs whose expression depends on virus origin and/or strain. Based on the antigenic properties of the viral surface glycoproteins, 18 HA (H1-18) and 11 NA (N1-N11) IAV subtypes have been identified in nature in multiple combinations. Except the H17N10 and H18N11 IAVs identified in fruit bat species of Guatemala and Peru^1,2^ respectively, the remaining IAV subtypes are found in wild aquatic birds of the order *Anseriformes* and *Charadriiformes*. Spread and diversity of IAVs across the globe depends on the migratory patterns of wild birds that include both regional and intercontinental movement^3^. From this primordial reservoir, IAVs with increased host range emerge and establish lineages in a wide range of avian and mammalian species. Most notably, the emergence of IAV in poultry, pigs, horses, and humans has been invariably associated with epidemic and/or pandemic episodes with significant loss of life and/or economic disruptions^4^. Based on the pathogenesis in chickens and/or presence of a polybasic amino acid cleavage site in the HA, IAVs that affect poultry are classified into two pathotypes as low pathogenicity avian influenza viruses (LPAIVs) and high pathogenicity avian influenza viruses (HPAIVs). Only viruses of the H5 and H7 subtype appear to be associated with either LPAIV or HPAIV strains, whereas other subtypes are typically characterized as LPAIVs^5^. The polybasic cleavage site in HPAIV enables the HA to be processed by endogenous cellular furin-like proteases enabling systemic infection. LPAIVs contain typically a monobasic cleavage site that limits HA processing to extracellular trypsin-like proteases, thus restraining the infection to mostly the intestinal and/or respiratory tract.

In Argentina, active influenza virus surveillance in wild birds was initiated in 2006^6^ as an integral part of worldwide efforts to better understand the ecology of these viruses and to better monitor the potential emergence of strains of concern for poultry, livestock, and humans. These studies led to the realization of IAVs in wild birds in Argentina have unique evolutionary patterns and the presence of a dominant gene segment constellation defined as South American lineage (SAm)^7^. IAVs of the H1, H4, H5, H7, H9, H10 and H13 subtypes were identified in Argentina, but the most common so far is the H6 subtype (∼41%) associated with the N2 subtype. Due to this observation and reports of H6 subtype IAV outbreaks in domestic poultry in Eurasia, North America, and Africa^8–11^, the prototypic influenza A/rosy-billed pochard/Argentina/CIP051-557/2007 (H6N2) strain, hereafter WT557/H6N2, was selected to establish its replication and pathogenic capacity in chickens. The resulting progeny was adapted through serial passage 20 times in chickens, and the resulting 20Ch557/H6N2 virus was also evaluated for replication and pathogenesis.

## RESULTS

### Serial passage in chickens improves replication of a wild bird origin H6N2 SAm IAV

We followed a strategy that was previously described for adaptation of a domestic duck origin H9N2 virus to Japanese quail and chickens using lung homogenates from the previous passage^12^. In this case, we attempted adaptation in chickens of a H6N2 IAV isolated from a rosy-billed pochard (WT557/H6N2). After 20 passages in chickens, the variant 20Ch557/H6N2 virus replicated more efficiently in chickens than the parental WT557/H6N2 strain (Fig 1). Overall, not only higher peak virus titers were observed but also the number of chickens and days with positive tracheal and cloacal swabs was higher in chickens inoculated with the 20Ch557/H6N2 virus compared to those inoculated with the WT557/H6N2 strain (Fig 1). Virus titers in trachea from 20Ch557/H6N2 inoculated chickens were significantly higher than WT557/H6N2 inoculated chickens at 1 and 3 dpi (Fig 1a, *p* = 0.0155 and *p* = 0.0093 respectively). Also, virus titers in cloaca from 20Ch557/H6N2 inoculated chickens were significantly higher than WT557/H6N2 inoculated chickens at 1 dpi (Fig 1b, *p* = 0.0372). Although overt signs of disease were not detected in neither group of the inoculated chickens, the results indicate that the chicken-adapted virus replicates better than WT557/H6N2 in trachea and cloaca, increasing viral shedding and for longer time post-infection; thus, our laboratory adaptation strategy has increased viral fitness of the duck origin SAm H6N2 virus in chickens.

**Fig 1.**
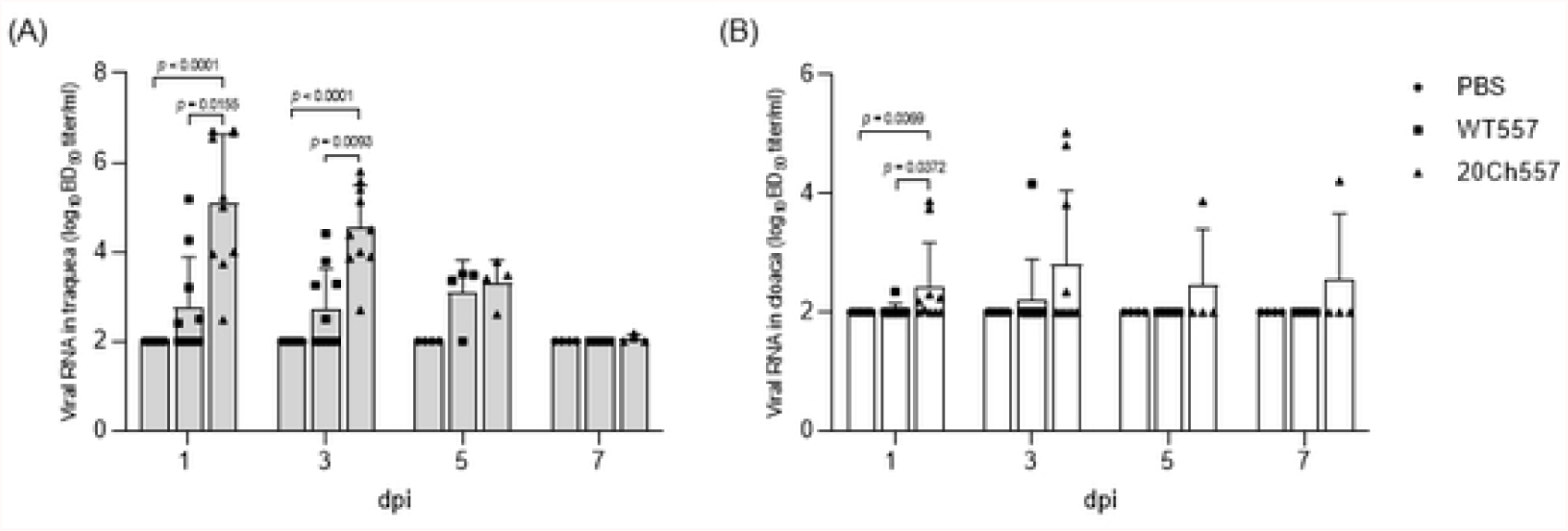
Replication of wild type and chicken-adapted H6N2 viruses in trachea and cloaca of chickens. Tracheal (A) and cloacal (B) swabs from control and IAV-infected chickens were analyzed for the presence of viral RNA at different days post-infection (dpi). For statistical purposes, tracheal and cloacal swabs without viral RNA detection were given a numeric value of 10^2^ EID_50_/ml, which represents the lowest detectable level of viral RNA with the RT-qPCR used. Graphs show the mean and standard deviation (SD). *p*-values were determined using ANOVA with multiple testing (Kruskall-Wallis test and Dunn’s Test).

### Differential replication of WT557/H6N2 and 20Ch557/H6N2 viruses in the tissues of chickens

To better determine the extent of virus replication of WT557/H6N2 and 20Ch557/H6N2 strains, a subset of chickens from each inoculated group (n=6 / group) was sacrificed at 3 dpi and bursa of Fabricius, lung, and intestinal tissues were collected. Mean bursa-to-body weight ratios were reduced in both IAV-inoculated groups in comparison to the negative control group (Fig 2, *p* = 0.0179 and *p* = 0.0039 for WT557/H6N2 and 20Ch557/H6N2 group respectively), consistent with IAV infection in chickens. All lung samples (n=6) from the 20Ch557/H6N2 inoculated group were RNA virus positive, in contrast to a single lung sample (n=1 out 6) from WT557/H6N2 inoculated group (*p* < 0.05) (Table 1). In addition, virus titers detected in lungs from the 20Ch557/H6N2 group were significantly higher compared to those from the WT557/H6N2 group (*p* < 0.05). No significant differences in neither number of positive samples nor in levels of virus titers in intestinal samples were observed for either IAV inoculated group. These results are consistent with the strategy of virus adaptation that selects for strains with improved respiratory tropism. In this case, such improvement does not appear to alter the natural tropism for the intestinal tract typical of wild aquatic bird origin IAVs.

**Table 1.** Replication of wild type and chicken-adapted H6N2 viruses in lung and intestine of infected chickens.

| Virus/Group | Viral RNA detection<sup>†</sup> (log<sub>10</sub> EID<sub>50</sub> titer/ml)<sup>ε</sup> |  |
| --- | --- | --- |
|  | Lung | Intestine |
| Control | 0/6 ($\leq 2$)<sup>A</sup> | 0/6 ($\leq 2$)<sup>A</sup> |
| WT557/H6N2 | 1/6<sup>a</sup> (2,1)<sup>A</sup> | 3/6<sup>a</sup> (2,66)<sup>A</sup> |
| 20Ch557/H6N2 | 6/6<sup>b</sup> (3,47)<sup>B</sup> | 4/6<sup>a</sup> (2,99)<sup>A</sup> |

<sup>†</sup> Number of positive chickens to IAV by RT-qPCR/total number of chickens.
<sup>ε</sup> For statistical purposes, lung and intestine without viral RNA detection were given a numeric value of 10<sup>2</sup> EID<sub>50</sub>/ml, which represents the lowest detectable level of viral RNA with the RT-qPCR used.

**Fig 2.**
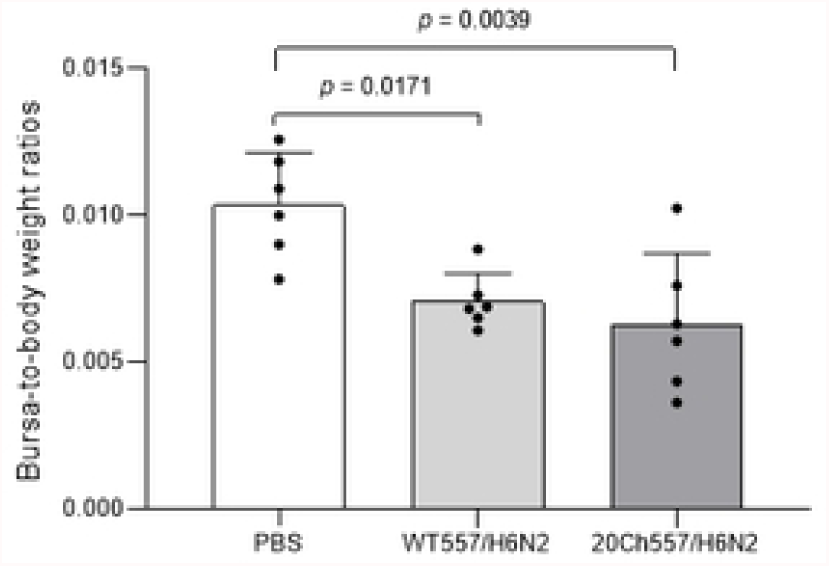
Tissue alteration after infection with wild type and chicken-adapted H6N2 viruses. Alterations of bursa-to-body weight ratios following infection with WT557/H6N2 or 20Ch557/H6N2 virus, 3 dpi. Mean values for IAV-infected and control animals were compared. The results are expressed as the mean (SD) from each group, n = 6. *p*-values were determined using ANOVA with multiple testing (Tukey’s Test).

Different superscript lowercase letters denote significant differences for number of positive chickens to IAV by RT-qPCR between groups; Chi-Square test, *p* < 0.05. Different superscript uppercase letters denote significant differences for mean viral titers between groups; Kruskal-Wallis test, *p* < 0.05.

Then, to determine whether SAm wild type or chicken-adapted H6N2 virus induces differences in the recruitment of mononuclear cells and/or lymphocyte subpopulations, the percentages of different lymphoid subsets in ileum from IAV-inoculated chickens were compared to those from control animals. Lamina propria and intraepithelial lymphocytic (IEL) cell compartments from ileum were analyzed. Remarkably, in both compartments, there was an increase of total mononuclear cells in response to the infection, being significantly higher only in the case of 20Ch557/H6N2 infected animals (Fig 3A, *p* < 0.05). Furthermore, an increase in the percentages of CD3+TCRγδ+ lymphocytes in the ileum of chickens inoculated with 20Ch557/H6N2 virus was observed (Fig 3B, *p* < 0.05), being in part due to an increase in CD3+TCRγδ+ CD8a+ subpopulation. Besides, macrophage population positive for staining of KUL01 marker showed a trend to increase in proportion in lamina propria from ileum from 20Ch557/H6N2 chickens (Fig 3C), as described during LPAIV infection^13^. The percentages of other subsets analyzed from ileum did not show differences between groups (data not shown). These results indicate that 20Ch557/H6N2 not only replicates better than WT557/H6N2 in the respiratory tract of chickens, but also induces a higher cellular immune recruitment in ileum.

**Fig 3.**
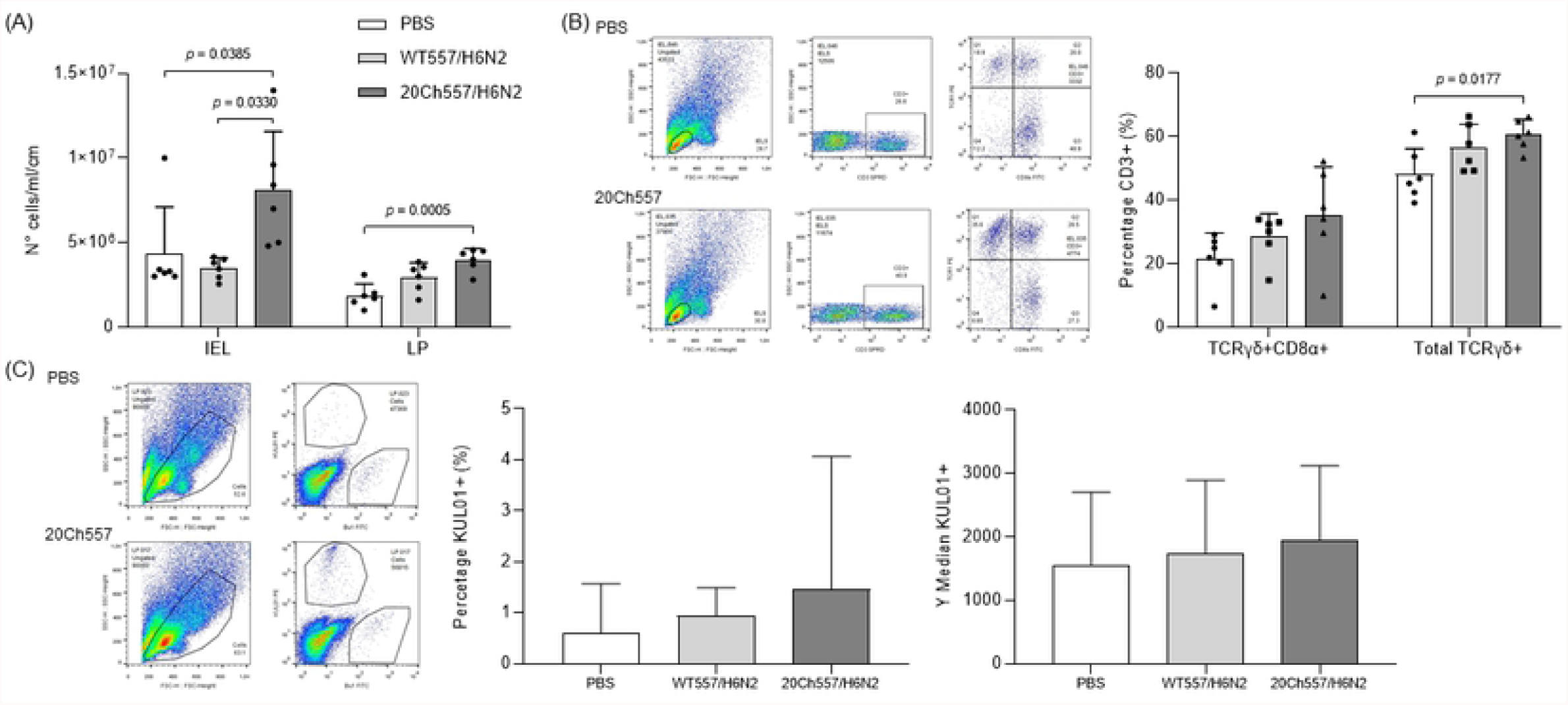
Differences in the recruitment of mononuclear cells and lymphocyte subpopulations in ileum of IAV-infected chickens. Six animals per group were euthanized at 3 dpi after infection with IAV or inoculation with PBS at 3 weeks of age. Ileum were collected and lamina propria and epithelial compartments were processed for flow cytometry analyses. (A) Mononuclear cell recovery from epithelial (IEL) and lamina propria (LP) compartment. Results are expressed as the mean (SD) of total amount of cells/ml recovered from each compartment from ten cm of ileum. p-values were determined using ANOVA with multiple testing (Dunn’s Test for IEL and Tukey’s Test for LP). (B) Chicken infection with 20Ch557/H6N2 virus increases TCRγδ+ IELs from ileum. Cells were labelled with anti-CD3, anti-CD8α and anti-TCRγδ antibodies conjugated to SPRD, FITC and PE respectively. Analysis was performed in the gate of lymphocytes according to forward/side scatter parameters, selecting CD3+ cells followed by TCRγδ-PE/CD8α-FICT. The results obtained in epithelial compartment analysis are shown, expressed as the mean in percentage (SD) from each group. p-values were determined using ANOVA with multiple testing (Tukey’s Test). (C) Macrophages in lamina propria from ileum of chickens infected with 20Ch557/H6N2 virus. Cells were labelled with anti-KUL01 and anti-Bu1 antibodies conjugated to PE and FITC respectively. Analysis was performed in the gate according to forward/side scatter parameters as shown followed by KUL01-PE/Bu1-FICT. The results obtained in lamina propria analysis are shown, expressed as the mean (SD) in percentage and MFI (SD) from KUL01+ population from each group.

### Mutations observed in 20Ch557/H6N2 virus genome after laboratory adaptation of WT557/H6N2 virus

To characterize the molecular features that allowed the chicken-adapted H6N2 virus to replicate more in trachea, lung, and cloaca of chickens (and also to a longer extent in trachea and cloaca), we compared deduced amino acid sequences from all open reading frames of WT557/H6N2 and 20Ch557/H6N2 viruses (Table 2).

**Table 2.** Comparison of amino acid changes in the proteins of WT557/H6N2 and 20Ch557/H6N2 viruses.

| Protein | Position | WT557/H6N2 | 20Ch557/H6N2 |
| --- | --- | --- | --- |
| PB2 | 748 | R | Q |
| PB1 | 484 | L | I |
| PB1-F2 | 83 | F | S |
| PA | 142 | K | R |
|  | 336 | R | L |
| PA-X | 142 | K | R |
| HA | 127 | V | I |
|  | 153 | Y | H |
|  | 182 | K | N |
|  | 346 | L | I |
| NA | 400 | N | S |
| M1 | 103 | L | I |
| NS1 | 86 | A | T |

Amino acid mutations were observed in the internal components of the chicken-adapted virus, either in structural or in nonstructural proteins. The only exceptions were NP and M2 proteins, which showed no amino acid substitutions. The PB2 protein of 20Ch557/H6N2 virus showed one amino acid difference compared to WT557/H6N2 virus at position 748 (R748Q). Another single amino acid substitution was observed in the PB1 protein of chicken-adapted H6N2 virus at position 484 (L484I). This mutation also produces the F83S change observed in the PB1-F2 protein of 20Ch557/H6N2 virus, which might be unique to the chicken-adapted virus since there are no PB1-F2 influenza sequences from SAm lineage in GenBank with this mutation. Then, there were two amino acid substitutions in the PA protein of 20Ch557/H6N2 virus, one at position 142 (K142R) - which is also encoded by PA-X protein of the adapted virus - and the other at position 336 (R336L). There were other two amino acid substitutions, one in M1 (L103I) and the other in NS1 (A86T) proteins of the 20Ch557/H6N2 virus.

We also found amino acid mutations in the superficial genes from 20Ch557/H6N2 virus (Table 2). The HA protein of chicken-adapted H6N2 has four amino acid changes at positions 127 (V127I), 153 (Y153H), 182 (K182N) and 346 (L346I) in the HA2 region of the Hemagglutinin. Finally, the NA protein of the 20Ch557/H6N2 virus has one mutation at positions 400 (N400S).

### Molecular changes in 20Ch557/H6N2 virus do not enhance transmission in chickens

To study direct contact transmission of WT557/H6N2 and 20Ch557/H6N2 viruses, four naïve chickens were placed together with infected birds at 1 dpi. Trachea and cloacal swabs were collected at different days post-contact (dpc) and sera collected at 20 dpc were tested by competitive enzyme-linked immunosorbent assay (ELISA) for specific antibodies against influenza A virus. Transmission studies indicated that the SAm wild type and chicken-adapted H6 viruses were not transmissible through direct contact. All swabs from direct contact animals were negative for the presence of viral RNA and blood samples did not show presence of antibodies against influenza virus (data not shown). When we analyzed serum samples from infected chickens, we observed 100% of infection with 20Ch557/H6N2 virus (4 out of 4 animals positive to ELISA) and 75% of infection with WT557/H6N2 virus (3 out of 4 animals positive to ELISA) (Table 3). Remarkably, any of the naïve animals developed antibodies against IAV along the study. These results show that the mutations obtained after serial lung passages of WT557/H6N2 virus in chickens improve virus’s infectivity, but do not enhance transmission in this host.

**Table 3.** Seroconversion of White Leghorn chickens against infection with wild type or chicken-adapted H6N2 virus.

| Virus/Group | Seroconversion<sup>†</sup> of birds |  |
| --- | --- | --- |
|  | Infected birds (21 dpi) | Direct contact birds (20 dpc) |
| WT557/H6N2 | 3/4 | 0/4 |
| 20Ch557/H6N2 | 4/4 | 0/4 |

dpi, days post-infection; dpc, days post-contact.
<sup>†</sup> Number of seropositive chickens by ELISA/total number of chickens.

### Antigenic relatedness between WT557/H6N2 and 20Ch557/H6N2 viruses

The antigenic properties of the SAm H6 viruses were investigated by using chicken antisera generated from the replication studies mentioned above. In general, antisera against the WT557/H6N2 and 20Ch557/H6N2 viruses showed low HI titers against the homologous virus (Table 4). It is notable that serum from chickens infected with 20Ch557/H6N2 showed lower HI titers against the homologous virus than against the heterologous virus. This result might be indicating that antigenicity of 20Ch557/H6N2 virus has been affected by the mutations introduced by the adaptation process, being the original WT557/H6N2 virus more immunogenic than the adapted one.

**Table 4.**
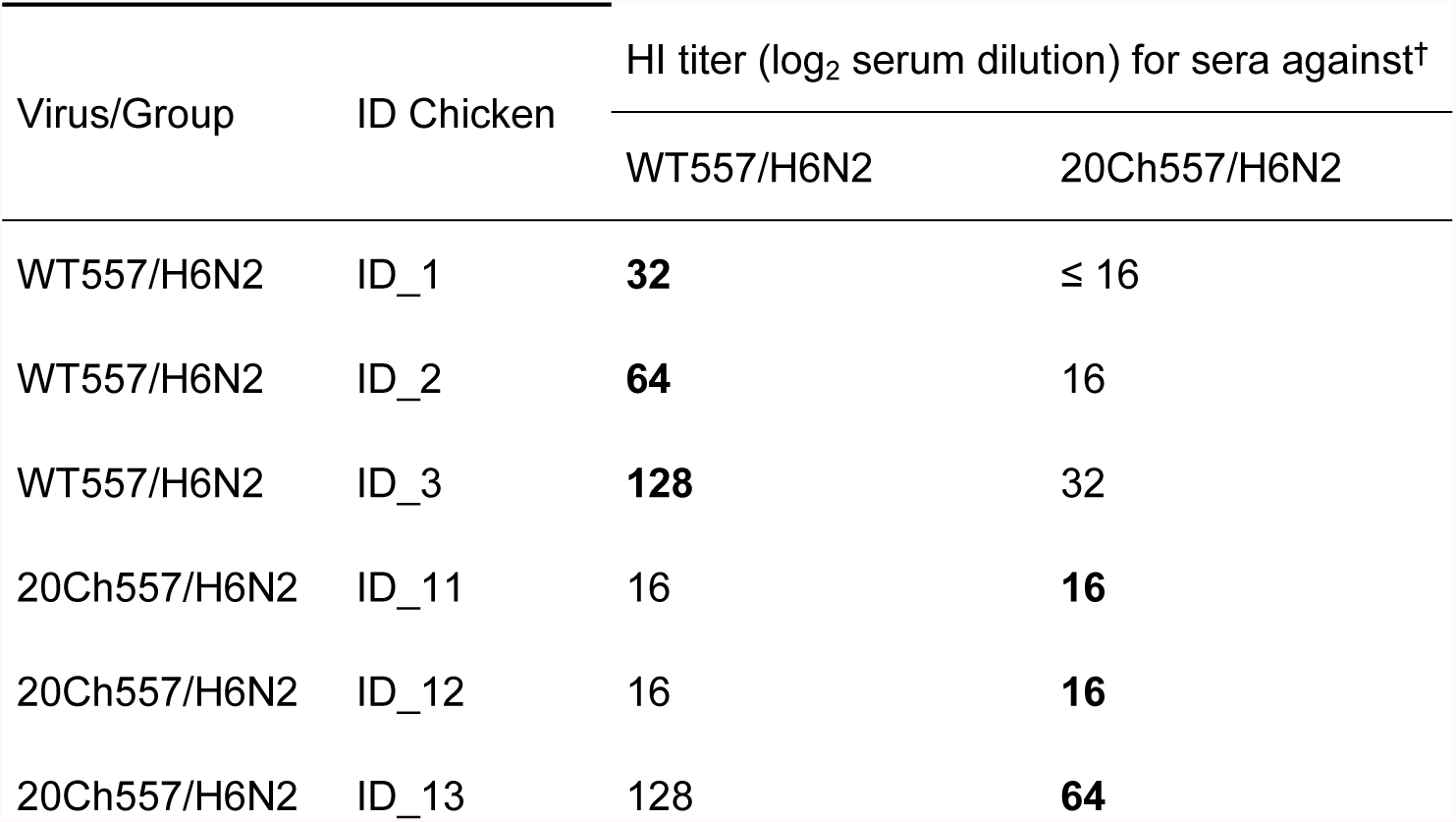
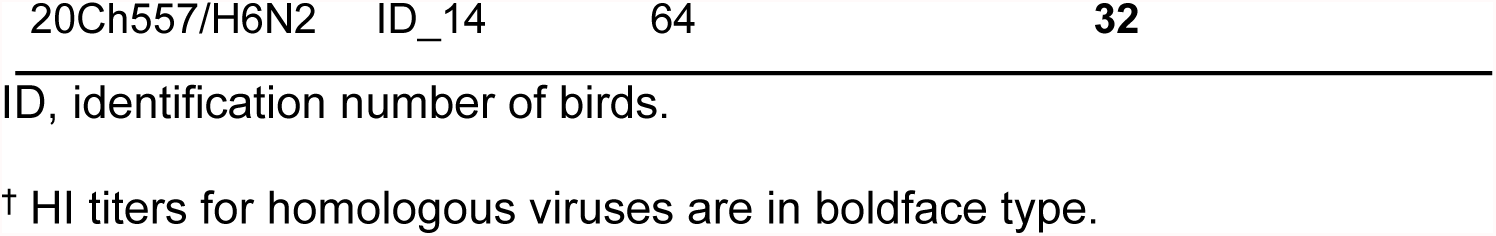
Serological data from chickens inoculated with WT557/H6N2 or 20Ch557/H6N2 virus at 21 dpi.

## DISCUSSION

Information on influenza A virus that circulates in wild birds in South America is scarce^14^. However, we have clear evidence of the circulation of a fixed SAm internal gene constellation in IAV isolated from wild birds in Argentina since 2006^7^. From the sixty IAV subtypes obtained to date - 22 IAVs already published and 38 IAVs detected from 2017 to 2019 - the predominant subtype is H6. Considering that Argentinean H6 IAVs could continue to evolve in wild birds, we conducted *in vivo* chicken studies with a H6 IAV of South American lineage isolated from wild duck and its adapted version in chickens, providing valuable and unknown knowledge on the pathogenesis and transmission of influenza A viruses isolated from wild birds in the region. Our results showed that, even though there was no evidence of clinical signs of disease compatible with IAV infection, both viruses infected chickens (observed mainly by seroconversion) but were not transmitted to other birds through direct contact (measured by virus detection from swabs and seroconversion). This, together with the significant bursa-to-body weight ratios reduction observed in both infected groups, demonstrated that WT557/H6N2 and 20Ch557/H6N2 viruses behave like LPAIV in chickens as observed with viruses from other lineages^15–17^. Our study showed that both viruses were detected in trachea and cloaca of infected chickens at 1 and 3 dpi, but only 20Ch557/H6N2 virus was detected in trachea and cloaca of infected chickens till day 7 post-infection. Also, viral titers were significantly higher in trachea and cloaca of 20Ch557/H6N2 infected chickens. When we analyzed the differential recruitment of mononuclear cells and lymphocyte subpopulations in the ileum from infected animals, we observed that 20Ch557/H6N2 chickens showed significantly higher percentage of CD3+TCRγδ+ lymphocytes in the epithelial cell compartment, indicating a difference in the recruitment of cells that might be implicated in the immunosurveillance against antigens in intestinal tract of chickens^18,19^. Together, these results evidence better replication and for longer time post-infection of 20Ch557/H6N2 virus with respect to the original virus obtained from ducks. Additional studies that analyze the lung immune response activated in chickens after infection with 20Ch557/H6N2 virus would shed more light in the virus-host interactions and the mechanisms involved in poultry to counteract an infection with SAm AIV strains more adapted to chickens.

Sequencing data from WT557/H6N2 and 20Ch557/H6N2 viruses was important to correlate the mutations observed (after 20 passages in chickens) to the higher viral replication observed in the respiratory tract of chickens infected with 20Ch557/H6N2 virus (Fig 1 and Table 1), which enabled the changing of viral tropism from exclusively gastrointestinal tract (known to occurred mostly in wild ducks) to a combination of respiratory and digestive tract (commonly observed in *Galliformes*)^20,21^. However, it is difficult to know if all or just only some of these amino acid changes are responsible for the higher fitness observed *in vivo*. The mutation observed in NS1 (A86T) protein of the 20Ch557/H6N2 virus has been already described in other influenza virus, however no clear association to species-specificity has been attributed to this variation^22,23^. Furthermore, no mammalian-associated virulence markers in PB2 (E627K and D701N)^24–26^ or NS1 protein (T92E)^27^ were found in the mutations observed in chicken-adapted H6N2 virus. Some authors studied the importance of four conserved PB1 motifs in influenza virus transcription/replication (Motif I to IV)^28^. For this, motif IV should have an isoleucine at position 484 in PB1, as we observed in 20Ch557/H6N2 virus. This mutation also produces the F83S change observed in the PB1-F2 protein of 20Ch557/H6N2. Considering that the PB1-F2 ORF from 20Ch557/H6N2 virus is 90 amino acids in length - as commonly found in AIV isolated from ducks - and that the all PB1-F2 proteins from AIV from SAm lineage have a phenylalanine at position 83 instead of serine, further functional analyses of the PB1-F2 protein from 20Ch557/H6N2 virus should be considered to determine its role in influenza pathogenicity in chickens.

With respect to the superficial genes, the HA protein of the 20Ch557/H6N2 virus differed from the WT557/H6N2 virus in the HA2 region at positions 127 (V127I), 153 (Y153H), 182 (K182N) and 346 (L346I). Interestingly, the three first amino acid substitutions in HA are part of the Receptor Binding Site (RBS). Functional studies are necessary to determine if these mutations increase the infective capacity of the adapted strain to chicken epithelial cells through a better interaction with sialic acid receptor. Considering that previous results showed that the interaction of recombinant HA proteins with sialic acid receptors may play a critical role in γδ T-cell activation^29^, together with our observation of an increment of this cell subpopulation in ileum of chickens infected with20Ch557/H6N2 virus, our results might be indicative of an important role in the mutations observed in the RBS of HA protein of the chicken-adapted strain. Furthermore, the mutation K182N incorporates a new glycosylation site in the RBS. Glycosylation in this region has been described as a mechanism used by influenza virus to escape from neutralizing antibody response^30,31^. Thus, this modification could also change antigenicity of the RBS and explain the lower antibody response elicited by the 20Ch557/H6N2 virus (Table 4) in spite of showing overall higher titers and longer period of viral detection compared to the parental strain. In addition, *Dong et al*. showed that the glycosylation of HA plays a significant role in the activation of γδ T cells^32^, suggesting that the increase in the percentage of CD3+TCRγδ+ lymphocytes observed in the epithelial cell compartment from our study could be related to the higher viral replication of 20Ch557/H6N2 virus and also to the acquisition of an additional glycosylation site in HA protein after chicken adaptation.

Then, the NA gene of the 20Ch557/H6N2 virus differed from the WT557/H6N2 virus at position 400 (N400S). This mutation in NA protein is not associated with drug resistance^33,34^. Interestingly, position 400 is in one of three loops that form a second sialic acid-binding site that is typical of bird-influenza virus^35,36^. Remarkably, the N in position 400 is mainly found in strains that infect different bird species outside *Galliformes* family, whereas S in position 400, as found in our chicken-adapted strain, is mainly found in viral variants identified in isolates from chickens^35,37^. This might be indicative that the N400S mutation observed in 20Ch557/H6N2 virus increases viral fitness in poultry. The available data of the mutations observed in IAV proteins, together with the results obtained from our chicken study, outlines the importance to determine which out of the thirteen amino acid changes found in 20Ch557/H6N2 virus are responsible for the higher fitness observed *in vivo*. Also, it would be interesting having knowledge of the number of passages that are necessary in chickens for the appearance of the critical mutations that change pathogenesis in this host.

In summary, our *in vivo* studies with IAV of SAm lineage showed that WT557/H6N2 and 20Ch557/H6N2 viruses replicated in SPF chickens but do not transmit by direct contact. The genetic modifications acquired by 20Ch557/H6N2 virus provided greater fitness in terms of infection, replication, and excretion in SPF chickens, triggering a greater immune response in the intestinal epithelium of this host at least at 3 dpi but showing a lower immunogenicity. Taken together, these results provide valuable information regarding the immunopathogenesis and transmission of avian influenza viruses from SAm lineage, which may be useful for future prevention strategies to improve the health status of poultry in the region. Finally, considering that IAVs of the H5, H7 and H9 subtypes have been found circulating in wild birds from South America, and that HPAIV outbreaks in poultry were reported in the region^38,39^; our results highlight the importance of further exploring the pathogenicity and transmission in chickens of other SAm IAVs to determinate their risk to poultry production.

## MATERIALS AND METHODS

### Viruses and inoculums

The A/rosy-billed pochard/Argentina/CIP051-557/2007 (H6N2) virus (WT557/H6N2) was isolated from waterfowl as described previously^40^. Stock of WT557/H6N2 virus was prepared by inoculation of embryonated chicken eggs. The chicken-adapted (20Ch557/H6N2) virus was isolated from the 20th chicken lung passage of the WT557/H6N2 virus using the methodology described by *Hossain et al*.^12^ Stock of 20Ch557/H6N2 virus was prepared by the inoculation of lung homogenates into the allantoic cavity of 10-day-old embryonated chicken eggs.

### Chickens and housing

Specific-pathogen-free (SPF) White Leghorn embryonated eggs (Rosenbusch S.A. CABA, Argentina) were purchased, incubated and hatched in an automatic incubator (Yonar, CABA, Argentina) under appropriate conditions, and randomly separated into groups of ten chickens. Groups were housed separately in sterilized isolators for chickens under negative pressure conditions (Allentown CH8ISOL) with food and water ad libitum throughout the experimental period. Animal care and experimental procedures were performed under ABSL3+ conditions with investigators wearing appropriate protective equipment and in accordance with the approved protocols N° 55/2013 of the National Institute of Agricultural Technology Ethics Committee (INTA, Argentina).

### Replication and transmission studies with H6N2 viruses

3-weeks old SPF White Leghorn chickens were used throughout the studies. Groups of chickens were inoculated intraocularly, intranasally, intratracheally and orally with WT557/H6N2 or 20Ch557/H6N2 virus containing 5×10^6^ EID_50_ in 1 ml or 1 ml PBS, respectively. At 1 dpi, four naive chickens were placed together with the infected birds to monitor transmission. Tracheal and cloacal swabs were collected at 1, 3, 5, 7 dpi in 1 ml freezing medium (50% glycerol in PBS containing 1% antibiotics) and stored at −80 °C until use. Virus titers in the swabs were determined by EID_50_ as previously described^41^. The limit of detection was determined to be 10^2^ EID_50_/ml; therefore, RT-qPCR negative samples were treated as ≤ 10^2^ EID_50_/ml. Chickens were observed daily for signs of disease (monitored for appetite, activity, fecal output, and signs of distress including cyanosis of the tongue or legs, ruffled feathers and respiratory distress), and serum samples were collected at 21 dpi.

To investigate the replication of WT557/H6N2 and 20Ch557/H6N2 viruses in the tissues of chickens (lung and intestine) 6 birds per group were anesthetized with isoflurane and euthanized using manual cervical dislocation at 3 dpi. Body and bursa of Fabricius were weighed and recorded in all animals.

### Epithelial cells isolation from ileum from chickens

Single cell suspensions of 10 centimeters of ileum were prepared as described previously^42^. Briefly, intestinal sections from the ileum were rinsed, cut longitudinally and washed with cold PBS containing 1% antibiotics. Following several washes, tissue sections were treated for 30 min at room temperature with PBS and 10 mM EDTA with continuous shaking. This was repeated and supernatants from both incubations were collected after centrifugation at 500 rpm for 3 min. Cells from supernatants were collected by centrifugation at 1200 rpm for 15 min and resuspended in RPMI 1640 medium. Finally, cells were passed through a 80 μm mesh (Cell Strainer, BD) and IELs were counted using trypan blue exclusion. For lamina propria compartment, ileum from chickens were cut mechanically and incubated with RPMI 1640 medium with Collagenase II in a concentration of 1 mg/mL (Gibco) for 40 minutes at 37 °C with shaker. Cells from supernatants were collected by centrifugation at 1200 rpm for 15 min and resuspended in RPMI 1640 medium. Finally, cells were passed through a 80 μm mesh (Cell Strainer, BD) and lymphocytes were counted using trypan blue exclusion.

### Flow cytometry analysis

Cells were diluted in staining buffer (PBS 1×, 2% FBS, 10 mM EDTA) and 1 × 10^6^ cells per well were seeded on 96 well-plates (V-shape) and washed twice with the same buffer. Staining was performed by resuspending the cellular pellet of each well with 100 μl of staining buffer including different combinations of antibodies, or as single-color staining for compensation. Cells were incubated at 4 °C for 20 min and washed twice with staining buffer by centrifugation at 1200 rpm for 3 min.

Monoclonal antibodies (mAbs) (CD3-SPRD, CD4-FITC, CD8α-FITC, CD8α-PE, TCRγδ-PE, BU1-FITC and KUL01-PE) were purchased from Southern Biotech. (Birmingham, AL), CD25-Alexa 647 was purchased from AbD Serotec (California, USA) and CD56-Alexa 647 was purchased from LSBio (LifeSpan BioSciences, Inc. Seattle, USA). Lymphocytes from lamina propria and epithelial cell compartments from ileum were stained with three mAb mix: CD3-SPRD, CD4-FITC, CD8α-PE and CD25-Alexa 647; CD3-SPRD, CD8α-FITC, TCRγδ-PE and CD25-Alexa 647; and CD3-SPRD, BU1-FITC, KUL01-PE and CD56-Alexa 647. All antibodies were titrated to determine the optimal staining concentration of each one.

Positive cells were analyzed with a FACS Calibur flow cytometer (BD Biosciences, San Jose, CA) and CellQuest software. Analyses were done on 20,000 events and discrete viable lymphoid cell populations were gated according to the forward/side scatter characteristics. Percentages of different lymphoid cell subpopulations in the ileum were determined by multiparametric analysis.

### Serology

Sera were tested by competitive enzyme-linked immunosorbent assay (ELISA) for specific antibodies against Influenza A Virus according to the manufacturer’s recommendations (IDEXX Laboratories Inc., Westbrook, ME) using 10-fold dilutions of each serum sample.

### HI Assay

Serum samples were collected to determine the antigenic relatedness between WT557/H6N2 and 20Ch557/H6N2 viruses at the end of the experiment. Sera were treated with receptor-destroying enzyme (Accurate Chemical and Scientific Corp., Westbury, NY, USA) to remove sialic acid receptors. The anti-viral antibody titers were evaluated using the HI assay system outlined by the WHO Manual on Animal Influenza Diagnosis and Surveillance (Webster et al., 2002). HI assays were performed using homologous and heterologous viruses.

### Amplification and sequencing of H6N2 viruses

Total RNA was extracted from stock virus in allantoic fluid using the QIAamp Viral RNA Mini Kit (Qiagen Inc., Valencia, CA, USA) according to manufacturer’s instructions. Reverse transcription and PCR amplification was performed using the universal primers described by *Hoffman et al*.^43^. PCR products were purified with a QIAQuick PCR purification kit (Qiagen) and sequencing was performed using the BigDye Terminator v3.1 Cycle Sequencing Kit on an ABI PRISM 3700 DNA Analyzer (Applied Biosystems) following the manufacturer’s instructions.

The consensus amino acid and nucleotide sequences for all eight gene segments of WT557/H6N2 and 20Ch557/H6N2 viruses were generated using BioEdit 7.

### Nucleotide sequence accession numbers

The genome sequences obtained in this study for WT557/H6N2 and 20Ch557/H6N2 stock viruses are available from GenBank under accession numbers CY067691 to CY067698 and ON306370 to ON306377.

### Statistical analysis

Data were tested for normal distribution (Shapiro-Wilk test) and homogeneity of variance (Leven’s test) prior to analysis using Statistical Package for the Social Sciences for Windows (SPSS, version 15.0, Chicago, IL, U.S.A.). An ANOVA test was used, and a Tukey’s test was applied when differences were detected. Variables that did not fit the ANOVA’s assumptions were analyzed by Kruskall-Wallis and Dunn’s Test. P-values of *p* < 0.05 were considered statistically significant.

Chi-Square tests were used to compare the number of animals that were positive to IAV between different experimental groups.

## ACKNOWLEDGMENTS

The authors thank to Mariela Gammella for her help with the animal studies. The National Institute of Allergy and Infectious Diseases (NIAID) Center for Research on Influenza Pathogenesis (CRIP) (contracts HHSN266200700010C and HHSN272201400008C) and the Instituto Nacional de Tecnología Agropecuaria (INTA) (PNSA 1115052 and PNSA 1115056) funded this study. This work was also supported by CONICET, through a fellowship gave to Agustina Rimondi during her postdoc. The content is solely the responsibility of the authors and does not represent official views of the National Institutes of Health.

## AUTHOR CONTRIBUTIONS

Author contributions were as follows: Study design: A.R., M.R., D.R.P. Virus sequencing: A.R., V.S.O. Animal studies with sample and data collection: A.R., V.S.O., I.S. Interpretation of protein mutations: A.R., G.P., M.R. Data analysis and interpretation of flow cytometry: A.R., I.S., M.R. Writing of the manuscript: A.R. Revision of the manuscript: M.R, D.R.P. All authors approved the manuscript before its submission.

